# Variation in symbiont density is linked to changes in constitutive immunity in the facultatively symbiotic coral, *Astrangia poculata*

**DOI:** 10.1101/2022.06.16.496460

**Authors:** Isabella Changsut, Haley R. Womack, Alicia Shickle, Koty H. Sharp, Lauren E. Fuess

## Abstract

Scleractinian corals are essential ecosystem engineers, forming the basis of coral reef ecosystems. However, these organisms are in decline globally, in part due to rising disease prevalence. Most corals are dependent on symbiotic interactions with single-celled algae from the family *Symbiodiniaceae* to meet their nutritional needs, however suppression of host immunity may be essential to this relationship. To explore immunological consequences of algal symbioses in scleractinian corals, we investigated constitutive immune activity in the facultatively symbiotic coral, *Astrangia poculata*. We compared immune metrics (melanin synthesis, antioxidant production, and antibacterial activity) between coral colonies of varying symbiont density. Symbiont density was positively correlated to both antioxidant activity and melanin concentration. Our results suggest that the relationship between algal symbiosis and host immunity may be more complex than originally hypothesized and highlight the need for nuanced approaches when considering these relationships.

## Introduction

Scleractinian corals are key ecosystem engineers, which create the structural basis of diverse coral reef systems [1]. However, the health of coral reefs worldwide is deteriorating, largely due to anthropogenic climate change [2]. Changing environmental conditions such as increased ocean temperatures and ocean acidification have led to coral die-offs [3]; global coral reef cover has declined by 50% from 1957 to 2007 [4]. The two largest drivers of coral mortality have been disease outbreaks and bleaching events [5-7]. Previous studies suggest extensive inter- and intraspecific variation in response to disease [8] and propensity to bleaching [9]. However, while the factors contributing to variation in bleaching susceptibility have been well studied in many coral species [9, 10], the mechanisms driving variation in coral disease susceptibility largely remain unknown.

The coral immune response consists of pathogen recognition, signaling pathways, and effector responses [11]. Corals have a variety of pathogen recognition molecules, such as Toll-like receptors and NOD-like receptors, capable of identifying a diversity of pathogens [12]. Post-recognition, signaling pathways appropriate defense mechanisms and trigger effector responses [12]. Corals use effector responses such as melanin production, antioxidants, and/or antimicrobial peptides to eliminate pathogens [12]. Preliminary evidence suggests that natural variation in several immune components might contribute to variation in disease resistance [13-15].

Beyond its role in pathogenic defense, the coral immune system also plays roles in the maintenance of symbioses [16, 17]. The onset and maintenance of coral symbiosis with Symbiodiniaceae is theorized to circumvent or modulate host immune response [18]. Furthermore, modification of immunity may extend beyond establishment of the relationship. In the threatened Caribbean coral *Orbicella faveolata* higher Symbiodiniaceae density was linked to negative effects on host immune gene expression [19]. Similarly, in *Exaiptasia diaphana*, symbiotic state was found to modulate NF-κB, a transcription factor responsible for numerous immune effector responses [20]. Still many questions remain regarding links between symbiosis and immunity in corals. To better understand how Symbiodiniaceae density and immunity might be linked in scleractinian corals, we investigated variation in constitutive immunity among colonies of the facultatively symbiotic scleractinian coral, *Astrangia poculata*, with variable symbiont densities.

## Materials & Methods

### Sample collection

*Astrangia poculata* colonies were collected from Fort Wetherill in Jamestown, Rhode Island in April 2021 (41°28′40″ N, 71°21′34″ W) at a depth of 10-15 meters, via SCUBA. Colonies were visually assessed and sorted into either high or low symbiont density groups (termed “brown” or “white” colonies respectively); 10 colonies of each type were collected. Visual assessment of colony color is a reliable method for distinguishing corals with high symbiont density (>10^6^ cells cm^-2^) from those with low symbiont density (10^4^-10^6^ cells cm^-2^ [21]). It should be noted that we use the terms “brown” and “white” as colonies grouped in the white category are rarely completely aposymbiotic. Following collection, the colonies were returned to Roger Williams University where they were maintained for several weeks in closed systems containing locally sourced seawater and fed three times weekly with frozen copepod feed. Samples were then flash frozen in liquid nitrogen and shipped to Texas State University for analyses.

### Protein extraction

Tissue was removed from colonies with extraction buffer (TRIS with DTT, pH 7.8) using protocols outlined by Fuess [22]. First, tissue was removed and isolated from a fixed surface area (2.14 cm^2^) on the flattest portion of the coral for Symbiodiniaceae density calculation. Then, tissue from the remaining fragment was removed and isolated into a separate aliquot. Both aliquots of tissue extracts were homogenized using a Fisherbrand Homogenizer 150 prior to downstream processing.

The Symbiodiniaceae aliquot was processed using a series of consecutive centrifugation and wash steps. The homogenate was centrifuged at 2000 RPM for 3 minutes and the supernatant was removed. The resultant pellet was resuspended in 500µL, and the product was centrifuged again using the same procedure. This step was repeated, and the sample was preserved in 500µL of 0.01% SDS in deionized water, stored at 4C.

The host aliquot was processed to obtain subsamples for protein activity assays and melanin concentration estimation. Following homogenization, 1 mL of the host aliquot was flash frozen, and stored at 20°C for melanin concentration estimation (see **Melanin** section). The remainder of the host aliquot was centrifuged for 5 minutes at 3500 RPM using an Eppendorf Centrifuge 5425 R. The resulting supernatant (protein enriched cell-free extract) was flash frozen in liquid nitrogen and stored at –80°C for downstream assays.

### Symbiont density

Symbiodiniaceae density was estimated using a standard hemocytometer and Nikon Eclipse E600 microscope. Symbiodiniaceae counts were repeated in triplicate and averaged to calculate symbiont density/tissue area.

### Biochemical Immune Assays

A Red660 assay (G Biosciences, St. Louis, Missouri) based on existing methods [23] was used to determine sample protein concentration and standardize assays. All immune assays were run in duplicates on 96 well plates using a Cytation 1 cell imaging multi-mode reader with Gen5 software (BioTek).

### Prophenoloxidase Cascade Assays (PPO, PO, and MEL)

Total phenoloxidase activity (PPO + PO) and melanin abundance was estimated using previously established methods [22] adapted to *A. poculata*. For total phenoloxidase activity, 20µL of coral extract were diluted into 20µL of 50 mM phosphate buffered saline (pH 7.0) in a 96 well plate (Greiner bio-one, Frickenhausen, Germany). Samples were incubated with 25µL of trypsin (0.1 mg/mL) for 30 minutes at room temperature, allowing for cleavage of PPO into PO. Post-incubation, 30µL of dopamine was added to each well. Absorbance was read every minute for 20 minutes at 490nm. Change of absorbance at the steepest point of the curve was used to calculate total phenoloxidase activity, standardized by protein concentration [22, 24].

To estimate melanin concentration, subsampled tissue extracts for the melanin assay were dried on a speed vac (Eppendorf, Vacufuge plus) in a tarred 1.5mL microcentrifuge tube. Dried tissues were weighed and processed to assess melanin concentration. Two hundred microliters of glass beads (10mm) were added to each tube. Samples were then vortexed for 10 seconds and 400uL of 10M NaOH was added to each tube. Tubes were vortexed for 20 seconds and incubated in the dark for 48 hours, with a second 20 second vortexing occurring after 24 hours. Post-incubation, the tubes were vortexed and then centrifuged at 1000 RPM for 10 minutes at room temperature. The resultant supernatant (40µL) was transferred to a ½ well UV plate (UV-STAR, Greiner bio-one, Frickenhausen, Germany). Absorbance was read at 410 and 490nm. We used a standard curve of melanin dissolved in 10M NaOH to calculate mg melanin from absorbance [22, 24]. Melanin concentration was standardized per mg of dried tissue weight.

### Antioxidant Assays

The activity of two coral antioxidants was investigated: catalase (CAT) and peroxidase (POX), following established methods [22, 24], adapted to *A. poculata*. Catalase was measured by diluting 5µL of sample with 45µL of 50mM PBS (pH 7.0) in a transparent UV 96-well plate (UV-STAR, Greiner bio-one, Frickenhausen, Germany). To initiate the reaction, 75µL of 25mM H_2_O_2_ was added to each well. Absorbance was read at 240nm every 30 seconds for 15 minutes. Scavenged H_2_O_2_ was calculated as the change in absorbance at the steepest portion of the curve. A standard curve was used to determine change in H_2_O_2_ concentration (mM), and results were standardized by protein concentration [22, 24].

To measure peroxidase activity, 20µL of sample was diluted in 20µL of 10mM PBS (pH 6.0) in a standard 96-well plate (Costar, Corning, Kennebunk, ME). Fifty microliters of 25mM guaiacol in 10mM of PBS (pH 6.0) was added to each well of the plate. To initiate the reaction, 20µL of 20mM H_2_O_2_ was added to each well. Absorbance was read every minute for 15 minutes at a wavelength of 470nm. Peroxidase activity was calculated as the change in absorbance at the steepest portion of the curve, standardized by protein concentration [22, 25].

### Antibacterial Assay

Antibacterial activity of *A. poculata* samples was assessed against *Vibrio coralliilyticus* (strain RE22Sm; provided by D. Nelson University of Rhode Island), a known coral pathogen [26]. Bacterial culture was revived from frozen stock and grown overnight in Luria broth (LB). After 24 hours, 1mL of bacterial culture was diluted in 100mL of mYP30 broth and grown for an additional 48 hours. Prior to assays, the culture was diluted to a final optical density at 600nm of 0.2. To initiate the assay, 140µL of bacterial culture and 60µL of sample, diluted to a standard protein concentration, were combined into wells of a sterile 96-well plate (Costar, Corning, Kennebunk, ME). Sample absorbance was read every 10 minutes at 600nm for 6 hours at 27°C. Change in absorbance during the logarithmic growth phase of the curve was used to calculate growth rate [22, 23].

### Statistical Analyses

Prior to statistical testing, outliers were identified and removed if necessary, using the ‘nooutlier’ function in R. Normality was also assessed, and data was transformed as needed; Symbiodiniaceae density was square root transformed. We assessed the effects of symbiont density on each of our immunological metrics using two approaches. First, we tested for differences in assay activity between colonies grouped as white or brown using a t-test. Second, we used a Pearson correlation test to assess direct correlations between symbiont density and assay activity. T-tests and correlations were run independently for each assay.

## Results

Statistical analysis revealed a significant association between symbiotic state and host immune phenotypes. Both melanin concentration (t-test, p=0.002; Figure 1A) and catalase activity (T-test, p=0.007; Figure1B) were significantly higher in brown colonies than white. Furthermore, melanin concentration (Pearson correlation, R=0.62, p=0.006; Figure 1C) and catalase activity (Pearson correlation, R= 0.62, p=0.007; Figure 1D) were significantly positively correlated to symbiont density. No other assays were significantly associated with symbiont state or symbiont density (**Tables 1-2**).

**Figure 1.**
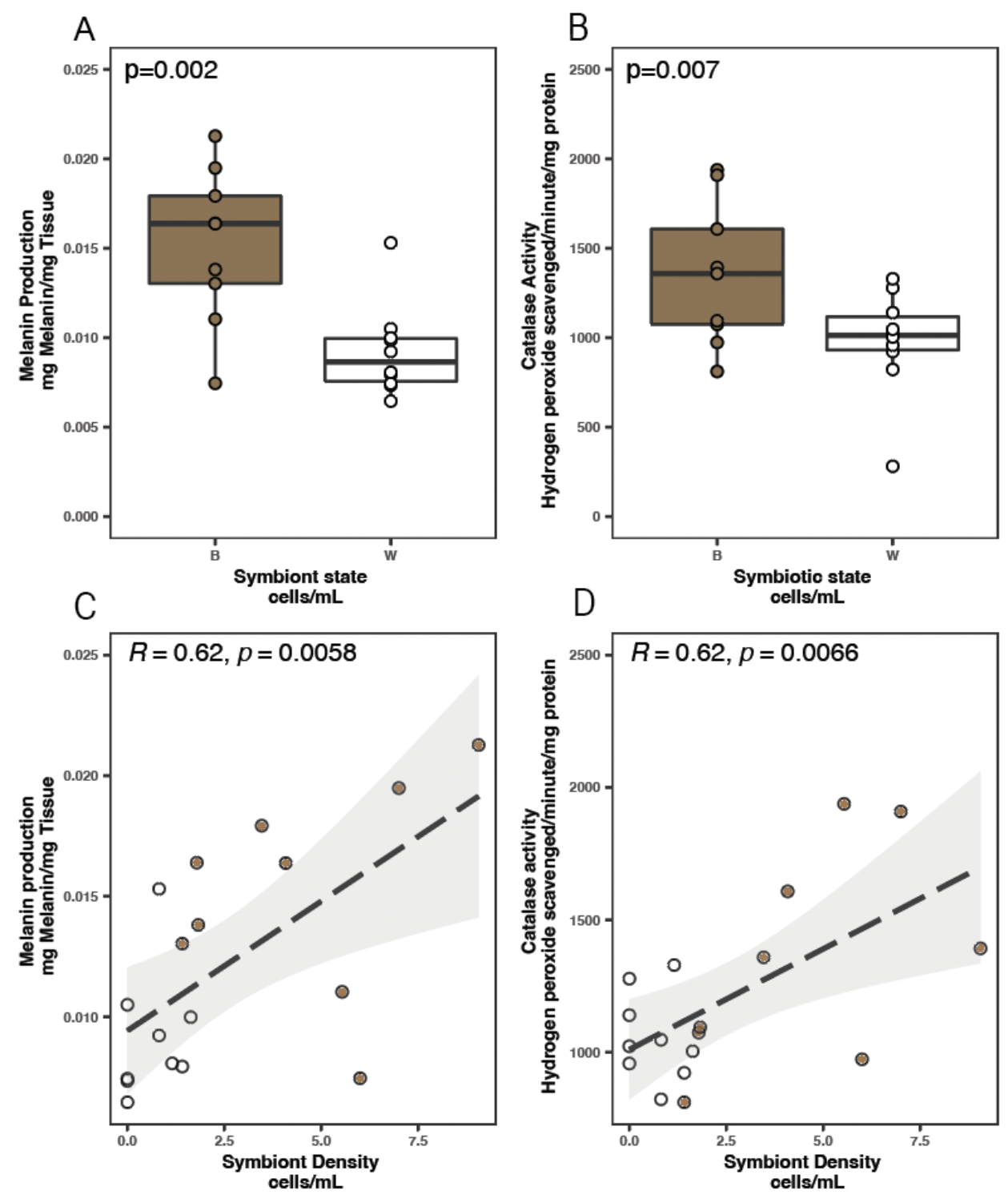
Both symbiont state and symbiont density affect melanin concentration and catalase activity. **A-B:** Box and whisker plots displaying differences in immune parameters between white and brown colonies for melanin **(A)** and catalase **(B). C-D:** symbiont-immune assay correlation results for melanin concentration (**C**) and catalase activity (**D)**.

**Table 1.** T-test results for each immunological assay.

| Assay | Statistic value | dF | p-value |
| --- | --- | --- | --- |
| Peroxidase | -0.591 | 14.4 | 0.564 |
| Prophenoloxidase | -0.865 | 17 | 0.399 |
| Catalase | 2.02 | 10.6 | 0.070* |
| Antibacterial | 1.55 | 17 | 0.139 |
| Melanin | 4.40 | 9.66 | 0.002* |

**Table 2.**
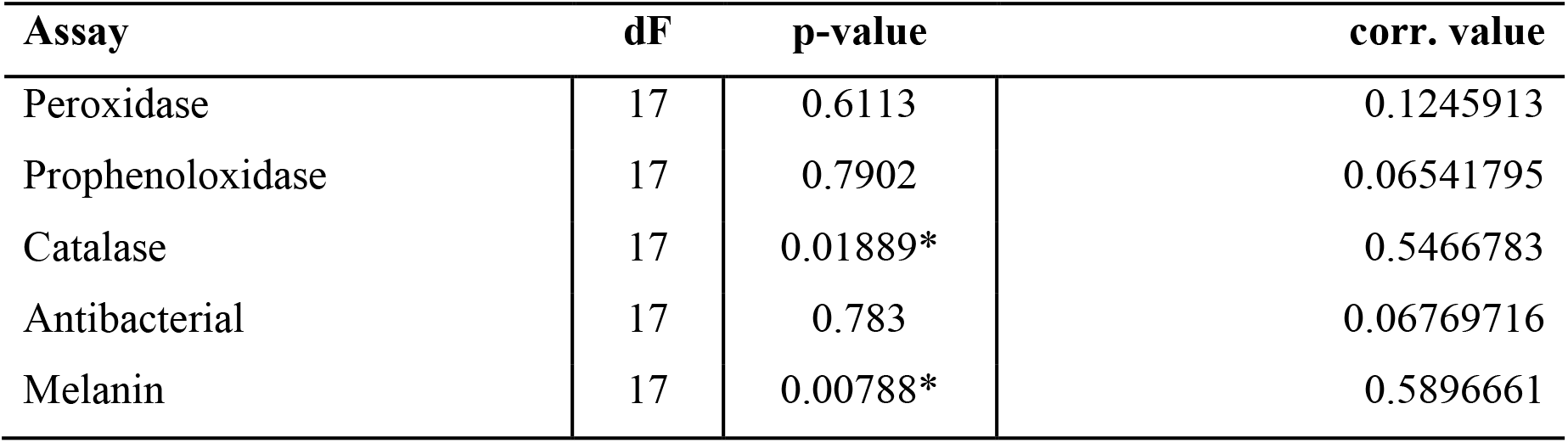
Pearson-correlation results between assay activity and square-root transformed symbiont density.

| Assay | dF | p-value | corr. value |
| --- | --- | --- | --- |
| Peroxidase | 17 | 0.6113 | 0.1245913 |
| Prophenoloxidase | 17 | 0.7902 | 0.06541795 |
| Catalase | 17 | 0.01889* | 0.5466783 |
| Antibacterial | 17 | 0.783 | 0.06769716 |
| Melanin | 17 | 0.00788* | 0.5896661 |

## Discussion

Here we used a facultatively symbiotic coral, *Astrangia poculata*, to investigate trade-offs between constitutive immunity and Symbiodineaceae density in corals. Past studies have suggested trade-offs between the maintenance of symbiotic relationship and immunity in obligately symbiotic corals [19]. In contrast, our results show no trade-offs between Symbiodineaceae abundance and constitutive immunity. Instead, we find a positive association between constitutive immunity and Symbiodineaceae density in *A. poculata*. These findings suggest that the relationship between algal symbiosis and immunity may be more complex than conventionally thought and highlight the need for further study of symbiosis-immune interplay in diverse systems.

Here we document positive correlations between symbiont density and two metrics of constitutive immunity: catalase activity and melanin concentration. Importantly, while both systems function in immunity, they also serve secondary roles in maintenance of coral-algal symbiosis [27]. While antioxidant activity is important in combating ROS bursts associated with pathogen defense, it is also important in general stress response, including response to thermal stressors [28]. Symbiont release of ROS is believed to be a cause of thermally induced bleaching, or breakdown of algal symbiosis [29]. Consistent with this theory, increased antioxidant production is associated with increased resistance to thermal bleaching [30]. Similarly, in addition to its roles in encapsulation of pathogens [12], melanin may play secondary roles in stress response, including protection of algal symbionts from UV damage (i.e., symbiont shading; [31]). Consequently, observed patterns of higher activity of these two pathways may be indicative of algal symbiont management and proactive stress mitigation mechanisms rather than direct consequences of symbiosis on immunity.

A second hypothesis could explain the observed associations between Symbiodineaceae density and immunity more generally: resource allocation theory. Resource allocation theory posits that organisms allocate a fixed energetic budget to competing needs (ex: growth, reproduction, and immunity; [32]). When energy budgets are fixed, increases in any one category come at the cost of another (i.e. tradeoffs; [32]). Consequently, energetic budgets can have significant impacts on resources allocated to immunity. For example, reductions in energy budgets caused by starvation resulted in decreased expression of immune genes and resistance to pathogens in the cnidarian *Nematostella vectensis* [33]. Indeed facultative symbiosis may be a natural source of variation in energetic budget; colonies of corals with variable densities of Symbiodineaceae may vary in their base energetic budget due to increased photosynthetically derived carbon. Past studies have linked increased photosynthetic energy acquisition to increased Symbiodiniaceae density [34, 35]. Consequently, increased densities of Symbiodiniaceae may increase a colonies total energetic budget, allowing for greater resource allocation to immunity and explaining positive correlations between certain immune phenotypes (catalase and melanin) and Symbiodiniaeae density.

In summary, our results highlight a positive association between Symbiodiniaceae density and immune parameters, which contrasts past studies of obligately symbiotic corals. This association is most likely either related to the dual function of these parameters or a consequence of increased energetic budgets associated with symbiosis. Importantly, our approach only measured a subset of potential effector responses. Future studies incorporating more responses or measures of receptor and signaling activity would improve interpretation of these trends. Additionally, our results are limited to the context of constitutive immunity; further studies considering pathogen response would be informative. Nevertheless, our data provides an important first step in highlighting the nuanced association between immunity and algal symbiosis in scleractinian corals.

## Acknowledgements

The authors extend appreciation to the annual Temperate Coral Research Conferences hosted by Roger Williams University, Boston University, and Southern Connecticut State University for fostering creative conversations and collaborations leading to this work. *Astrangia poculata* colonies were collected by Michael Lombardi, Ocean Opportunity, LLC.

## Funding

This work was supported by Texas State University (startup funds to LEF) and the NIH NIGMS Institutional Development Award (IDeA) Network for Biomedical Research Excellence (grant P20GM103430 to KS).

## Author Contributions

IC, KS, & LEF designed the experiment. KS & AS planned and executed sample collection and shipping. IC, HW, & LEF processed samples. IC & LEF conducted statistical analyses. All authors contributed to manuscript writing and revision.

## Notes

### Competing Interest Statement

The authors have declared no competing interest.

